# Development of novel high-affinity nanobodies against EGFR for cancer therapy

**DOI:** 10.1101/2025.03.16.643542

**Authors:** Mauro Heitrich, Marisa M. Fernandez, Diana C. Aguilar-Cortes, Santiago Werbajh, Gabriela Canziani, Vanesa Zylberman, Luisina B. Ripari, Guillermo A. Videla-Richardson, Emilio L. Malchiodi, Osvaldo L. Podhajcer, Sabrina E. Vinzon

## Abstract

The epidermal growth factor receptor (EGFR) is a member of a family of transmembrane tyrosine kinase receptors that plays a pivotal role in regulating diverse cellular processes such as cell proliferation, survival, migration, and differentiation. Aberrant activation of EGFR signaling has been implicated in various pathological conditions, particularly cancer, making it an attractive target for therapeutic intervention. While several anti-EGFR monoclonal antibodies have been developed and demonstrated their clinical value for the treatment of various solid tumors, smaller antibody fragments such as nanobodies (Nb) offer distinct advantages over conventional antibodies, including reduced immunogenicity and enhanced tumor penetration. In this paper, we report the isolation and characterization of two novel high-affinity Nb targeting EGFR. These Nb were identified and characterized using ELISA, flow cytometry, microscopy, and SPR. Furthermore, these Nb and bivalent Nb engineered from them were tested for their effects on cancer cell proliferation. We demonstrate that the novel Nb exhibit high affinity and potent anti-tumor activity *in vitro* in their bivalent form, positioning them as promising candidates for cancer treatment.

## 1. Introduction

The epidermal growth factor receptor (EGFR) is one of the four members of the HER family of transmembrane tyrosine kinase receptors, which includes EGFR/HER1/erbB1, HER2/erbB2, HER3/erbB3, and HER4/erbB4 (Tebbutt et al., 2013). The EGFR signaling pathway plays a pivotal role in regulating diverse cellular processes such as proliferation, survival, migration, and differentiation. Aberrant activation of EGFR signaling has been implicated in various pathological conditions, particularly several types of tumors (Kumagai et al., 2021), and increased levels of EGFR are associated with advanced disease stages and poor prognosis (Krasinskas, 2011; X. Liu et al., 2017). Consequently, EGFR is considered an attractive target for cancer therapy, leading to the development and investigation of numerous pharmacological approaches (Lee, 2017). Small molecule tyrosine kinase inhibitors (TKIs), such as gefitinib, lapatinib, afatinib and erlotinib, have been widely used in clinical practice, and are effective against various cancers, including lung, breast, prostate and colorectal cancers (Huang et al., 2020). However, due to the emergence of resistance, most patients eventually experience disease progression (X. Liu et al., 2017). Another targeted therapy to inhibit EGFR signaling involves the use of monoclonal antibodies (mAbs), which have demonstrated efficacy in specific cancer types harboring EGFR alterations (Huang et al., 2009). Notably, cetuximab and panitumumab, two well-known EGFR-directed mAbs, have shown clinical benefits by inhibiting ligand binding, receptor dimerization, and downstream signaling, ultimately leading to cell-cycle arrest and apoptosis (London & Gallo, 2020). However, mAbs used in cancer therapy are not without disadvantages. Mouse-derived mAbs can lead to rapid clearance from the system, reduced efficacy, or hypersensitivity reactions in patients, while humanized antibodies only partially reduce immunogenicity (Chen et al., 2023; Hong & Sloane, 2019; Isabwe et al., 2018; Khoo et al., 2017). Tumors often develop resistance to cetuximab and panitumumab due to mutations in the EGFR extracellular domain, which impair antibody binding and lead to clinical relapse (Cai et al., 2020). Furthermore, the large molecular weight of mAbs reduce their penetration into solid tumors, and their production in mammalian cells is costly (Xenaki et al., 2017). Consequently, smaller antibody fragments are becoming a real alternative to conventional mAbs for cancer therapy.

Nanobodies (Nb), derived from heavy-chain-only antibodies in camelids and sharks, offer a promising avenue for cancer therapy and fighting resistance (Iezzi et al., 2018; Rahbarizadeh et al., 2011). With their nanoscale dimensions, they can penetrate deep into tumors and even cross the blood-brain barrier, potentially being more effective in targeting EGFR-overexpressing cells (Santos et al., 2021; Tsitokana et al., 2023). Moreover, they can be engineered to have high specificity and affinity for EGFR while minimizing off-target accumulation, potentially overcoming some resistance mechanisms seen with mAbs (Muyldermans et al., 1994; Santos et al., 2021). They possess a high refolding capacity and can tolerate extreme conditions such as high temperatures, elevated pressures, and non-physiological pH levels (Kunz et al., 2018). Due to their inherent modularity, nanobodies offer advantages over conventional antibody fragments when used as part of more intricate constructs (Iezzi et al., 2018; Yang & Shah, 2020). Additionally, their simple and cost-effective production using microbial systems, along with their lack of post-translational modifications, yields homogeneous products (Muyldermans et al., 1994). Recently, several anti-EGFR nanobodies have been developed and used for diagnosis and therapy of EGFR overexpressing tumors (Sharifi et al., 2021). Notably, these small binding moieties, such as the nanobody 7D12, can target specific epitopes overlapping with the EGF-binding site on EGFR (Gainkam et al., 2008; Schmitz et al., 2013; Tintelnot et al., 2019). When combined with other single-domain antibodies targeting different epitopes, such as 9G8 which binds to the interphase between domain II and III (Schmitz et al., 2013), the resulting bivalent Nb is more effective in EGFR downregulation (Heukers et al., 2013; Roovers et al., 2011).

In this paper, we employed a combination of immunization and panning strategies to isolate novel nanobodies targeting EGFR in cancer cells (for a schematic representation of the overall strategy, see Fig. 1). We immunized llamas with a combination of cells and viral-like particles (VLP) overexpressing EGFR and screened the phage displayed-libraries with both VLPs and the EGFR ectodomain (ECD) fused to the Fc portion of IgG. Two potent anti-EGFR Nb were identified and characterized by ELISA, flow cytometry, microscopy and surface plasmon resonance (SPR). Additionally, these Nb as well as bivalent Nb engineered from them, were tested for their effects on the proliferation of EGFR-overexpressing cancer cell lines. The novel Nb showed high affinity and potent anti-tumor activity *in vitro* in their bivalent form, indicating their potential as candidates for cancer treatment.

**Figure 1.**
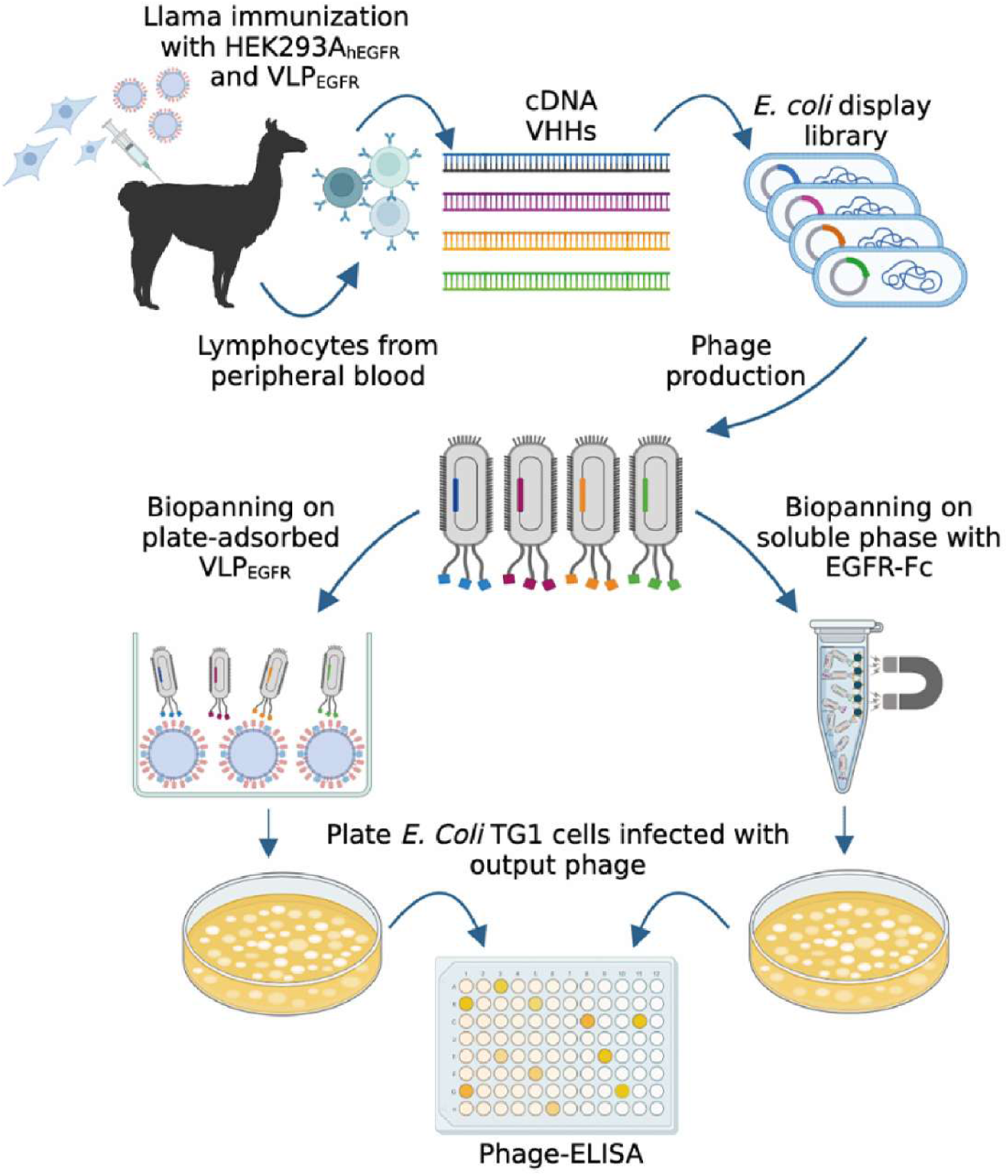
Schematic representation of the generation and display of the anti-EGFR VHH immune library on *M13 bacteriophage* and the methodology used for selection of anti-EGFR Nb.

## 2. Results

### 2.1. Different strategies for selection of EGFR-specific clones from the phage display library produce distinct outcomes

To generate a cell line with high EGFR expression, HEK293A cells were transfected with a plasmid encoding EGFR, and a stable clone, HEK293A_hEGFR_, was selected as described in the methods section. EGFR expression in HEK293A_hEGFR_ cells was found to be over 600 times higher than in HEK293A cells (Fig. S1). In developing EGFR-specific nanobodies (Nb), we implemented a dual-phase immunization strategy designed to enhance target specificity and immune response effectiveness. First, llamas were immunized with HEK293A_hEGFR_ cells which express high levels of EGFR, to elicit a strong antibody response against the target. A late-stage boost was administered with Virus-Like Particles (VLPs) displaying EGFR to preferentially expand clones recognizing EGFR epitopes. The induction of a specific humoral immune response was confirmed by testing the animals’ sera by ELISA before and after immunization. A specific response towards EGFR was induced in one llama (Fig. S2A), although the elicited antibodies were also directed towards antigens other than EGFR, as evidenced by the reactivity of immune sera against VLP_null_ (Fig. S2B). The antibody titers were sufficiently high to proceed with library construction, which contained approximately 10^7^ colony-forming units (CFU), with 86.7% of full-size inserts according to colony PCR (Fig. S3A).

To select for high-affinity EGFR-specific nanobodies, we applied two different panning strategies using either plate-adsorbed VLPs or purified EGFR ectodomain (EGFR-ECD). VLP-based panning allowed us to maintain the native-like presentation of EGFR, exposing phages to conformational epitopes similar to those encountered *in vivo*. Meanwhile, panning on purified EGFR-ECD, free from background proteins, offered a highly specific approach by limiting non-EGFR antigens, thereby reducing the likelihood of enriching irrelevant clones. This dual approach aimed to isolate high-affinity specific clones through either strategy, leveraging their distinct advantages to explore complementary avenues and increase the overall likelihood of success.

The first selection attempt was performed on plate-adsorbed VLPs, following standard biopanning methods (Pardon et al., 2014). Screening of 92 random clones from round 3 of this panning by VLP-ELISA resulted in only 9 clones with modest VLP_EGFR_/VLP_null_ selectivity ratios (SR >1.3). Sequencing revealed that 7 of these 9 clones contained aberrant Nb sequences, with most showing different VHH fragments connected head to tail through the PstI restriction site. The remaining 2 clones failed to show EGFR specificity upon further testing. Sequencing of 12 random isolates from the original library revealed each to be unique, but two clones contained aberrant sequences, either head-to-tail tandem VHHs or VHH fragments. These aberrant sequences do not yield functional antibodies due to premature stop codons introduced by the antisense orientation of the upstream fragment. The enrichment of such clones likely resulted from their growth advantage during amplification steps rather than specific binding to the target antigen. To reduce the spurious amplification of aberrant clones, we decided to tightly control the steps before each round of panning. We found that an amplification condition of 4 hours at 28 °C was optimal to amplify phages to a level sufficient to serve as input for the next round without over-amplification. Additionally, we monitored the quality of each input/output by colony PCR. PCR analysis showed an in-frame rate of 90-100% for the libraries after each round (Fig. S3). Using this strategy, we performed 3 rounds of affinity selection by phage display on purified EGFR-ECD or VLPs. Ninety-two phages from the third round of each selection were individually tested by VLP-ELISA for specific binding to VLP_EGFR_ and non-specific binding to VLP_null_. Eleven clones from the VLP-based panning and 26 clones from the EGFR-ECD panning were positive in the initial screening (i.e., they displayed OD_450_ >1.0, with >3-fold higher signal on target VLP_EGFR_ over VLP_null_, Fig. 2). These clones were further characterized by EGFR-ECD ELISA, which showed that all clones with SR <7 that passed the initial criteria had reactivities to EGFR-Fc close to background levels (corrected OD_450_ <0.060). Of the clones obtained after screening the VLP-panned output, only 3 out of 11 showed binding to EGFR-ECD. A total of 29 positive colonies with a binding ratio of more than 7 were further sequenced, revealing 12 different sequences. Finally, the EGFR-specific Nb were classified into 11 families based on the diversity of their CDR3 amino acid sequences (Table 1).

**Figure 2:**
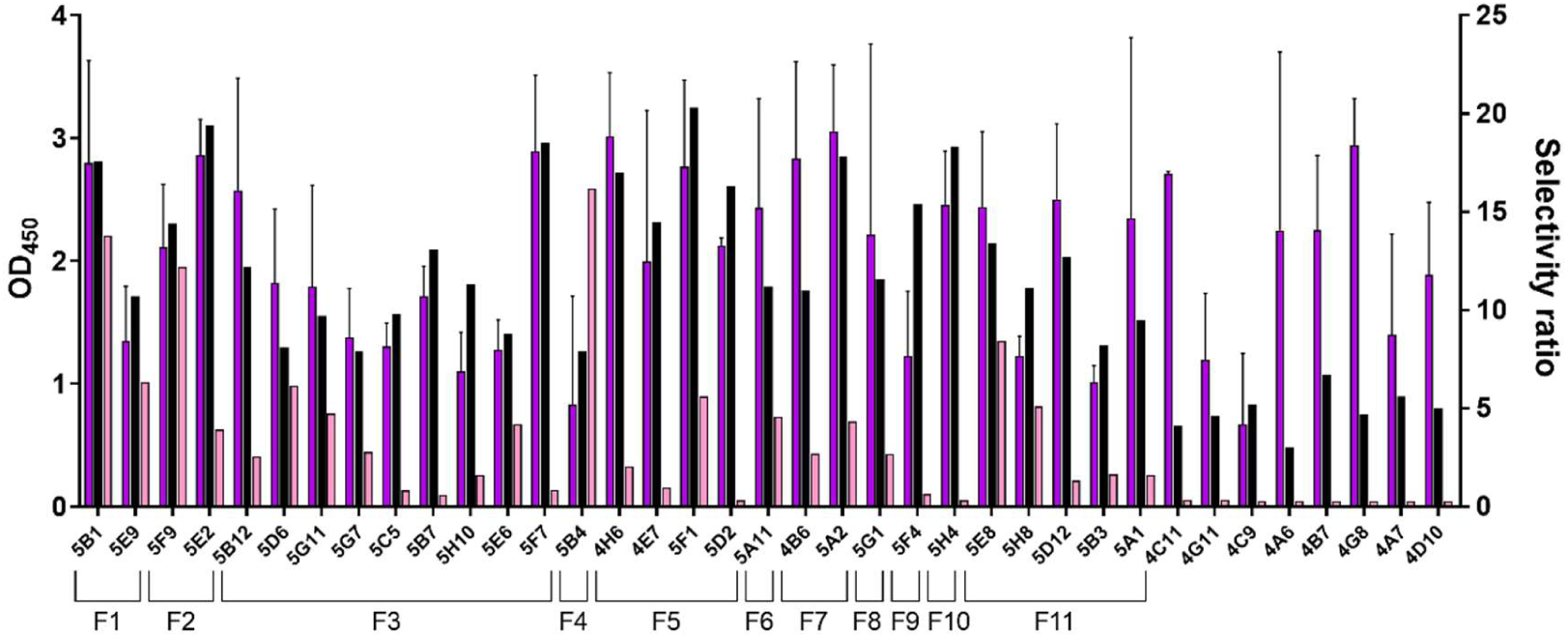
Screening of selected phages by ELISA. Phages from the third round of each selection strategy were evaluated by VLP-ELISA. Purple bars show the OD_450_ against VLP_EGFR_. Black bars depict the selectivity ratio (OD_450_ against VLP_EGFR_ / OD_450_ against VLP_null_). Pink bars indicate the OD_450_ against EGFR-ECD. Only the 37 clones which displayed an OD_450_ against VLP_EGFR_ >1.0, with >3-fold higher signal on target VLPs over VLP_null_ are shown. Clones whose name starts with 4 are from the VLP panning strategy, while those starting with 5 are from the EGFR-Fc/magnetic beads panning strategy.

**Table 1:**
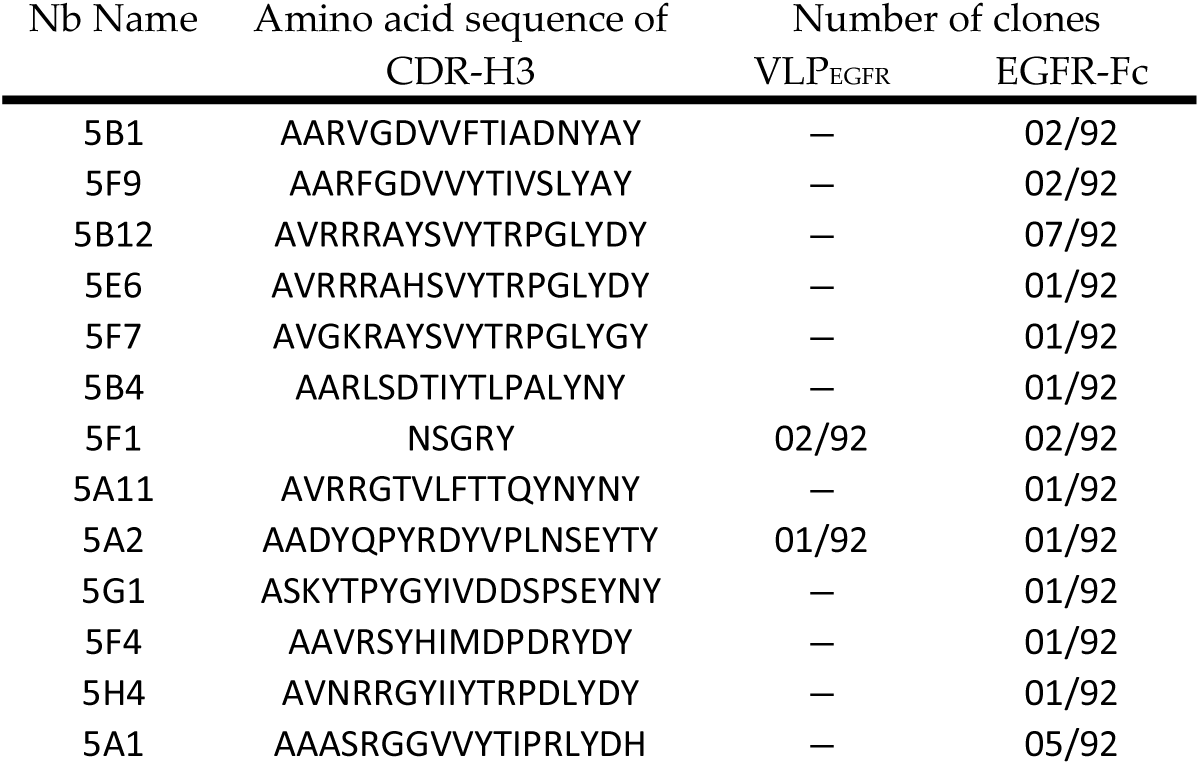
Nbs against EGFR selected with VLP_EGFR_ and EGFR-Fc.

### 2.2. Selection of Nanobodies 5B1 and 5B12

To determine the binding properties of the selected VHHs, one sequence from each Nb family was expressed as a His-tagged VHH in the periplasm of *E. coli* WK6. Seven unique Nbs were successfully obtained (Fig. S4A), and the purified Nb were tested for their ability to specifically bind EGFR-expressing cells by FACS (Fig. 3A). Flow cytometry analyses revealed that two Nbs, 5B1 and 5B12 (sequences in Fig. S5), specifically bound to EGFR^hi^ A431 cells. Two additional Nbs, 5A2 and 5F4, also bound to EGFR^hi^ A431 cells but exhibited lower affinity (Fig. 3A). In contrast, three of the tested Nbs, 5A1, 5B4 and 5G1, showed no differential binding between EGFR^hi^ A431 cells and EGFR^lo^ MCF7 cells (Fig. 3A). Binding of a control Nb (anti-BLS) to EGFR^hi^ A431 cells was at background levels. Based on these results, 5B1 and 5B12 were selected for further characterization. Preparations of 5B1 and 5B12 were analyzed for their aggregation status by size-exclusion chromatography (SEC), confirming monodisperse peaks corresponding to the molecular weight of the VHH monomer (Fig. S4C). For comparison, the public-domain VHH 7D12, known for its binding to EGFR, was included as a control. Its aggregation status similarly showed a single monodisperse SEC peak (Fig. S4C).

**Figure 3:**
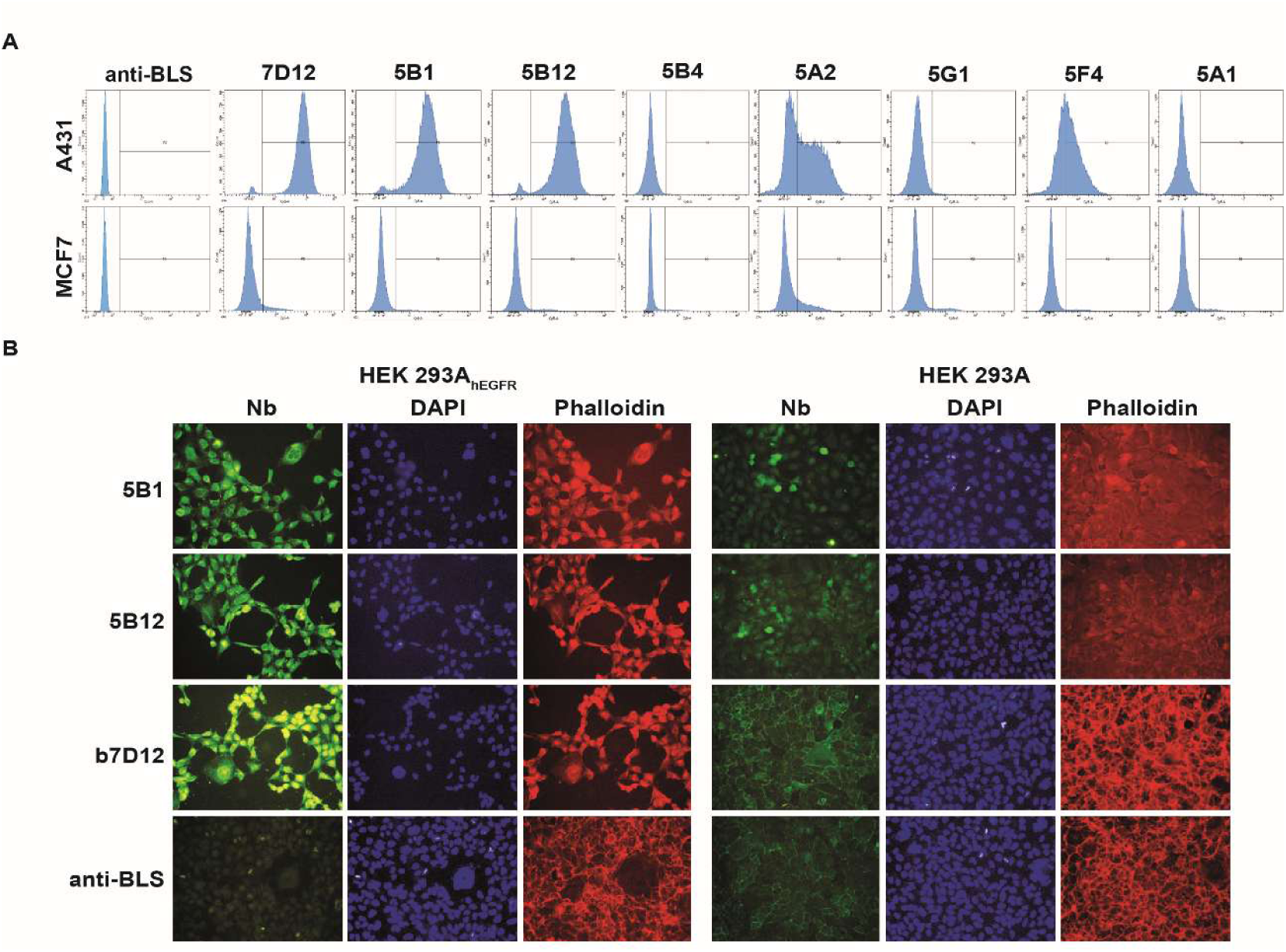
5B1 and 5B12 specifically bind EGFR-expressing cells. Binding analysis by flow cytometry. Purified samples of each Nb were assayed at 500nM by FACS against EGFR^hi^ A431 and EGFR^lo^ MCF7 cells. (B) Specific staining of EGFR-overexpressing HEK293A cells with the selected Nb. Immunofluorescence images show HEK293A_hEGFR_ and HEK293A cells stained with the selected Nb against EGFR. Nb binding was detected using anti-His Tag mAb followed by anti-mouse IgG (green). Cells were counterstained with DAPI (blue) to visualize the nucleus and phalloidin (red) to reveal the F-actin cytoskeleton.

The selected Nbs were further tested for their ability to specifically bind HEK293A_hEGFR_ cells by immunofluorescence microscopy (Fig. 3B). Cells were stained with 5B1, 5B12, bivalent 7D12 (b7D12, positive control) or anti-BLS Nb (negative control). Both 5B1 and 5B12 specifically stained HEK293A_hEGFR_ cells, with strong signals consistent with high EGFR expression levels. In HEK293A cells, staining by 5B1 and 5B12 was observed at background levels, similar to that seen with 7D12. This low-level staining aligns with the basal expression of EGFR in HEK293A cells, which can account for some degree of reactivity.

### 2.3. 5B1 and 5B12 bind to EGFR with high affinities

Binding of 5B1 and 5B12 to EGFR was further characterized by determining their half maximal effective concentration (EC_50_) on A431 cells using flow cytometry (Fig. 4). Both nanobodies, along with the gold standard 7D12, bound to A431 cells with nanomolar affinities, with the novel Nbs displaying an even stronger binding to EGFR overexpressing cells compared to 7D12 (5B12= 4.7 nM; 5B1= 76.9 nM; 7D12= 146.8 nM).

**Figure 4:**
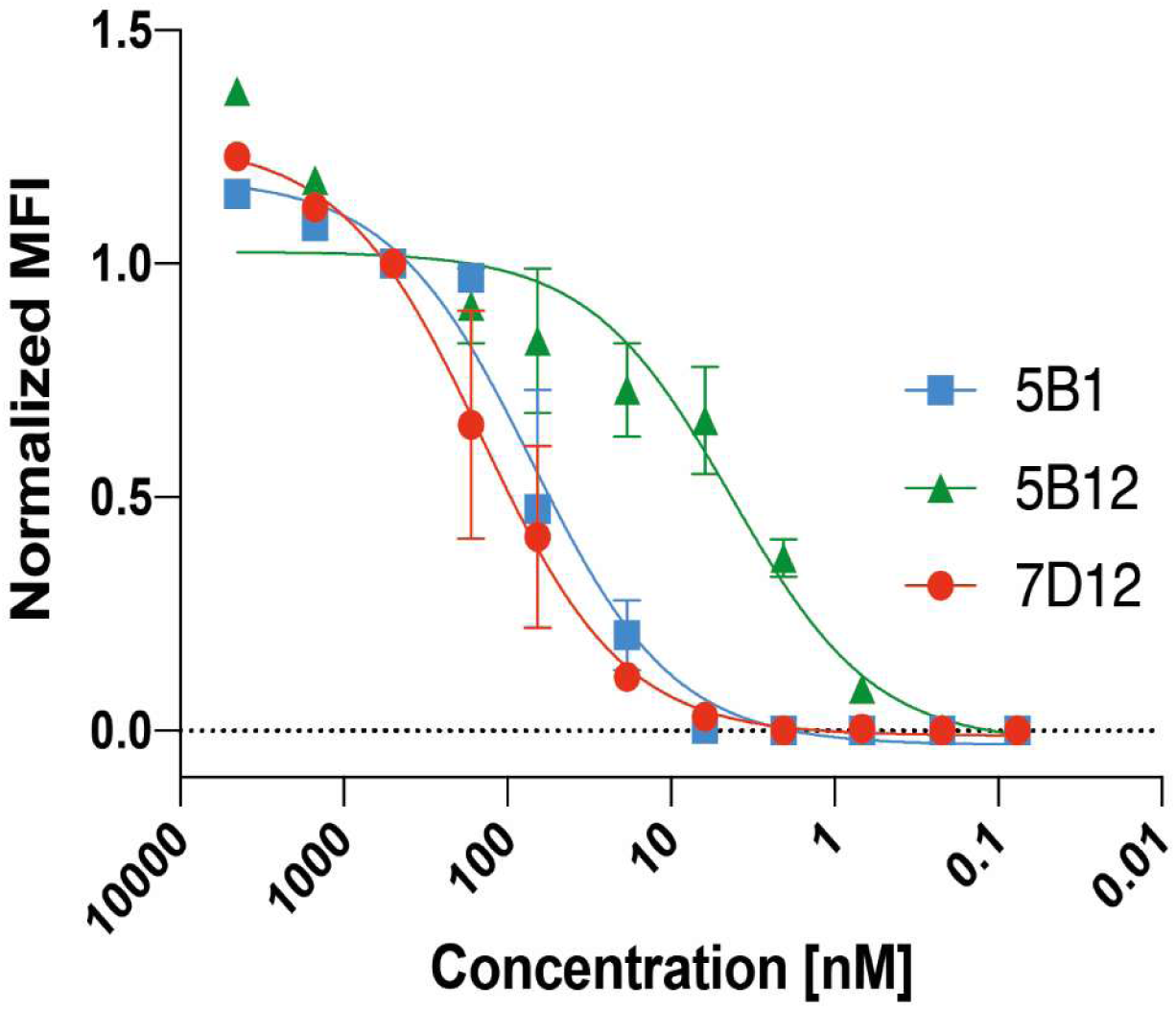
Saturation binding apparent K_D_ and B_max_ values for Nb on A431 cells, as determined by flow cytometry. Binding curves of median fluorescence intensity (MFI) versus antibody concentration are shown. Binding at each concentration was normalized to the MFI at 500 nM for each Nb. MFIs were fitted using nonlinear regression curves in GraphPad Prism 8.

The kinetic binding properties of the VHH monomers were confirmed on the purified EGFR ectodomain by Surface Plasmon Resonance (SPR) at a flow rate of 50 µl/min to reduce mass transport effects and minimize re-binding (Table S1 and Fig. S6). While the *K_D_* values of 5B1 and 5B12 were comparable, their kinetic profiles differed markedly. 5B12 exhibited a significantly faster association rate (*k_on_*) and a higher dissociation rate (*k_off_*) compared to 5B1, potentially conferring distinct biological behaviors to these nanobodies (Table 2). Although both nanobodies displayed slightly lower K_D_ values than 7D12, these differences are attributed primarily to their distinct kinetic parameters rather than a significant increase in affinity (Table 2). Notably, the lower *k_off_* of 5B1 suggests longer-lived complexes with EGFR, whereas the higher *k_on_* of 5B12 could enhance its capacity for rapid rebinding in vitro.

**Table 2:**
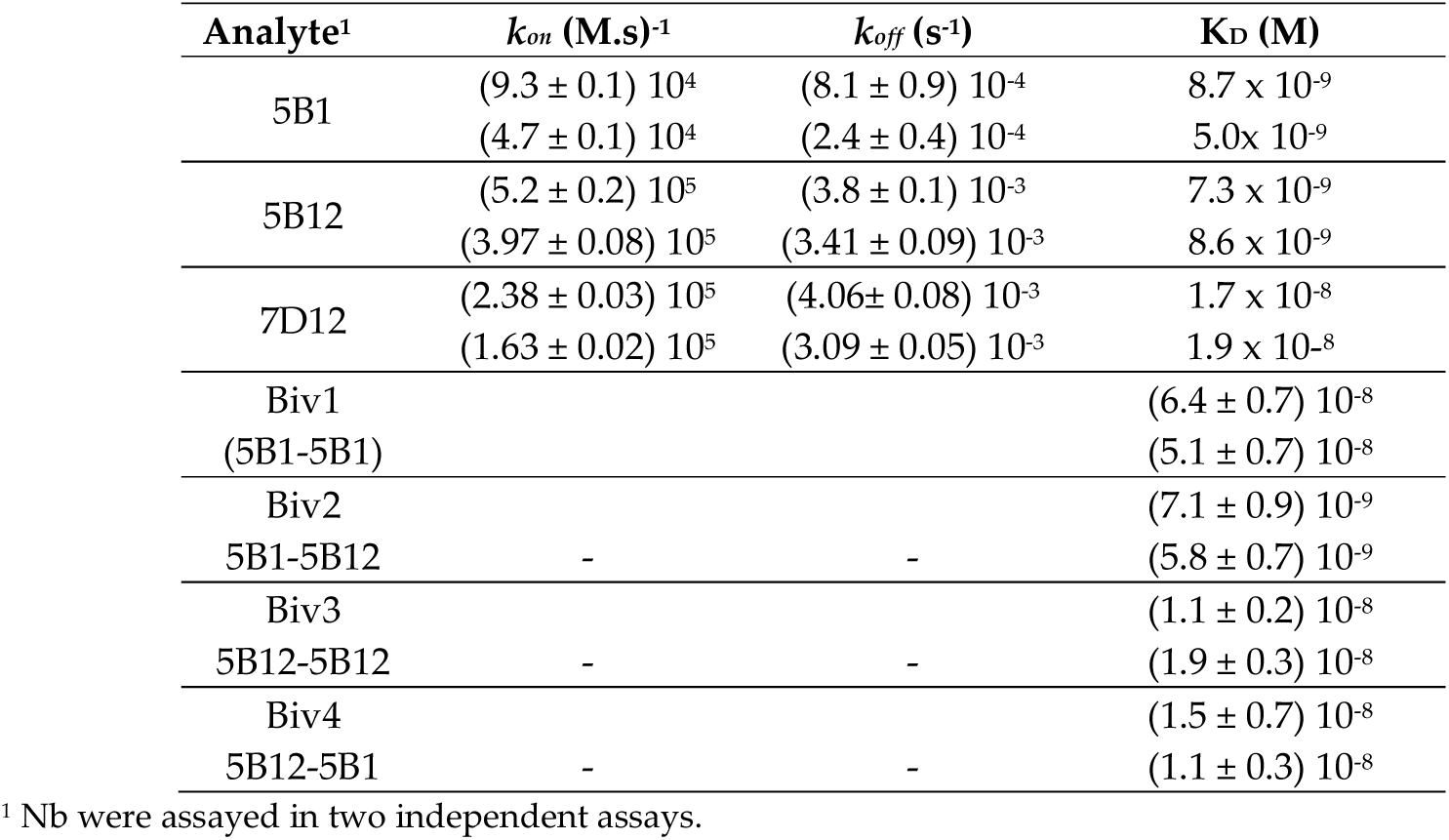
Kinetic parameters of the interaction between EGFR and VHHs determined at a flow rate of 50 µl/min.

| Analyte<sup>1</sup> | $k_{on}$ (M.s)<sup>-1</sup> | $k_{off}$ (s<sup>-1</sup>) | $K_D$ (M) |
| --- | --- | --- | --- |
| 5B1 | $(9.3 \pm 0.1) 10^4$ | $(8.1 \pm 0.9) 10^{-4}$ | $8.7 \times 10^{-9}$ |
| | $(4.7 \pm 0.1) 10^4$ | $(2.4 \pm 0.4) 10^{-4}$ | $5.0 \times 10^{-9}$ |
| 5B12 | $(5.2 \pm 0.2) 10^5$ | $(3.8 \pm 0.1) 10^{-3}$ | $7.3 \times 10^{-9}$ |
| | $(3.97 \pm 0.08) 10^5$ | $(3.41 \pm 0.09) 10^{-3}$ | $8.6 \times 10^{-9}$ |
| 7D12 | $(2.38 \pm 0.03) 10^5$ | $(4.06 \pm 0.08) 10^{-3}$ | $1.7 \times 10^{-8}$ |
| | $(1.63 \pm 0.02) 10^5$ | $(3.09 \pm 0.05) 10^{-3}$ | $1.9 \times 10^{-8}$ |
| Biv1 | | | $(6.4 \pm 0.7) 10^{-8}$ |
| (5B1-5B1) | | | $(5.1 \pm 0.7) 10^{-8}$ |
| Biv2 | | | $(7.1 \pm 0.9) 10^{-9}$ |
| 5B1-5B12 | - | - | $(5.8 \pm 0.7) 10^{-9}$ |
| Biv3 | | | $(1.1 \pm 0.2) 10^{-8}$ |
| 5B12-5B12 | - | - | $(1.9 \pm 0.3) 10^{-8}$ |
| Biv4 | | | $(1.5 \pm 0.7) 10^{-8}$ |
| 5B12-5B1 | - | - | $(1.1 \pm 0.3) 10^{-8}$ |

<sup>1</sup> Nb were assayed in two independent assays.

### 2.4. 5B1 and 5B12 target different epitopes and can be engineered as biparatopic nanobodies

Competition ELISA was performed to gain insight into the epitopes targeted by the novel Nbs. Phages displaying the 5B1, 5B12 or the control 7D12 Nbs were incubated with VLP_EGFR_ in the presence of increasing concentrations of cetuximab, which targets a well-characterized epitope overlapping the domain III ligand-binding region of EGFR, also recognized by 7D12 (Schmitz et al., 2013). The 5B1 phage exhibited a competition profile similar to 7D12, indicating at least partial overlap of their epitopes (Fig. 5). In contrast, the 5B12 phage was only partially displaced by cetuximab, suggesting that it targets a distinct epitope compared to 5B1.

**Figure 5:**
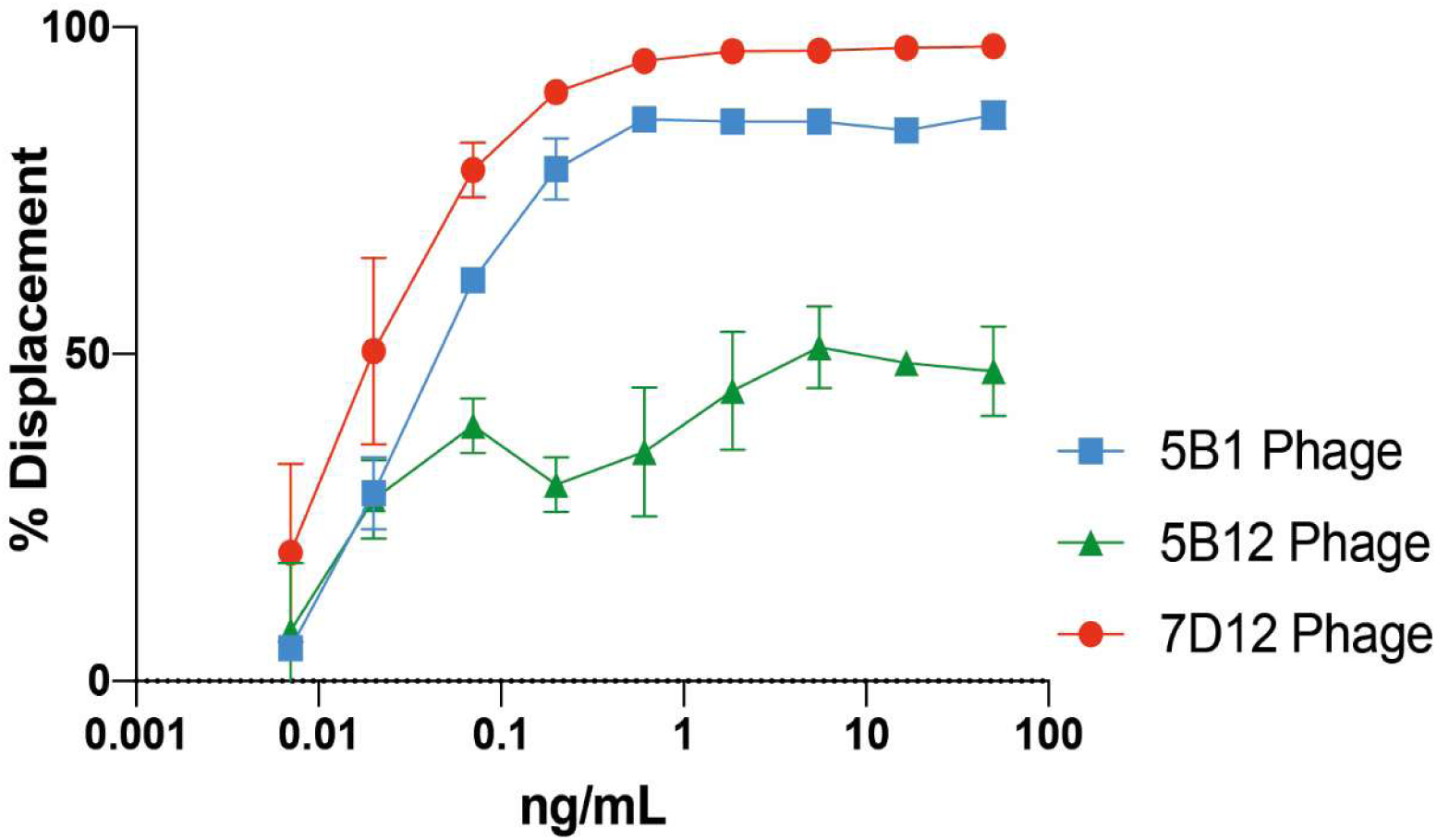
Characterization of VHH binding sites. EGFR-specific VHH-phages were tested in competition ELISA using VLP_EGFR_ as the antigen. Competition is expressed as the percentage of displacement by increasing concentrations of cetuximab.

### 2.5. Bivalent nanobodies engineered from 5B1 and 5B12 inhibit tumor cell proliferation

The effects of the selected nanobodies on cell proliferation were assessed using EGFR-overexpressing A431 cells. Monovalent nanobodies, including 5B1, 5B12, and the control 7D12, did not significantly affect cell proliferation, in contrast to cetuximab, which strongly reduced cell viability (data not shown). Cetuximab’s efficacy is attributable to its bivalence, a feature previously shown to be critical for effective EGFR blockade by antibodies (Fan et al., 1994; Harwood et al., 2017). To explore the functional potential of bivalence, we engineered bivalent nanobodies (bNb) by covalently linking two nanobodies with a flexible peptide linker in tandem orientation.. Four bNb variants (5B1-5B1, 5B1-5B12, 5B12-5B1 and 5B12-5B12) were expressed in *E. coli* WK6 (Fig. S4B). Biparatopic constructs 5B1-5B12 and 5B12-5B1 were specifically designed to investigate the impact of VHH placement on the N-terminal versus C-terminal positions on their properties (Beirnaert et al., 2017; Walter et al., 2022). Despite their modified composition, none of the bivalent constructs exhibited higher affinities than their monovalent nanobodies (Table 2 and Fig. S7).

*In vitro* studies revealed that all bNb variants displayed significant anti-proliferative effects on A431 cells, achieving maximal inhibition rates comparable to cetuximab, albeit with weaker inhibition at lower concentrations (Fig. 6A). These findings align with their ability to block EGF-induced EGFR phosphorylation. EGFR phosphorylation status was assessed by stimulating A431 cells with human EGF; cetuximab served as a positive control for blocking EGF binding, and untreated cells served as the negative control. Consistent with their anti-proliferative effects, only bivalent constructs (Biv1, Biv3 and Biv4) significantly reduced pEGFR levels (Fig. 6B), suggesting that bivalence is crucial for both receptor blockade and functional inhibition.

**Figure 6:**
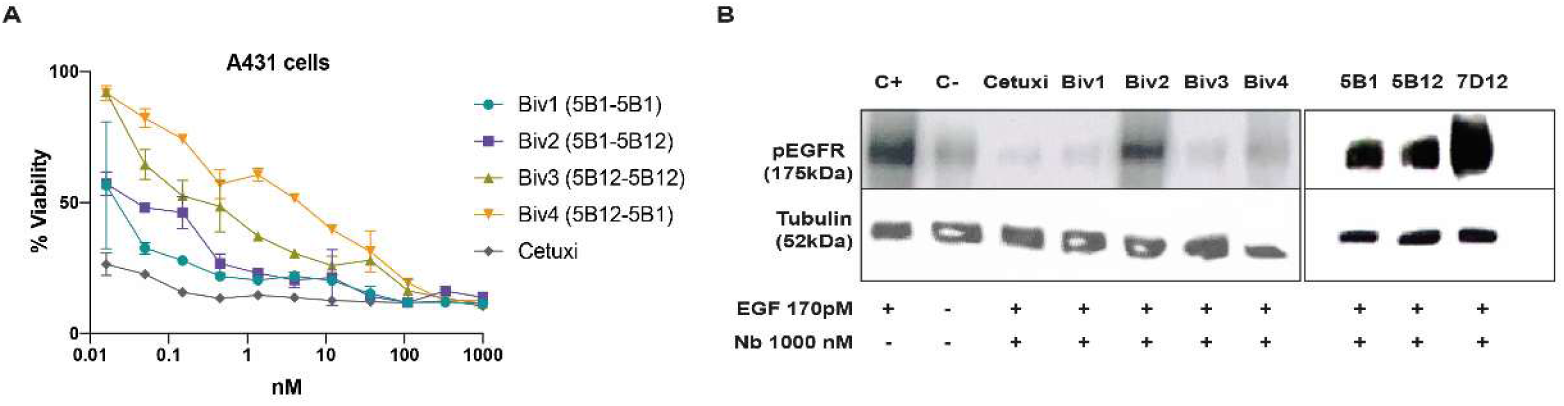
Bivalent nanobodies inhibit A431 proliferation. (A) EGF-dependent A431 cells were treated with different equimolar concentrations of the bivalent nanobodies or cetuximab, and the inhibition of cell proliferation was measured in triplicates after 6 days by MTS assay. (B) Phosphorylated EGFR status in A431 cells pre-incubated with serial dilutions of the mono/bivalent nanobodies or cetuximab was determined by Western Blotting after treatment with 170 pM of EGF for 10 min.

To further evaluate the therapeutic potential of these nanobodies, we tested their capacity to inhibit the *in vitro* proliferation of glioma stem cell-enriched cell lines expressing or not the truncated EGFR variant mutation vIII (Videla Richardson et al., 2016). Interestingly, the biparatopic Nbs 5B1-5B12 and 5B12-5B1 inhibited the proliferation of G05 cells expressing the mutated EGFRvIII; of note, 5B12-5B1 achieved up to 30% inhibition at 10 nM (Fig. 7). Surprisingly, none of the bNbs affected the G08 cell line expressing wild-type EGFR (Fig. 7). This differential effect was also observed with the commercial mAb cetuximab (Fig. 7), likely indicating a shared target-specific mechanism between the bivalent constructs and cetuximab.

**Figure 7:**
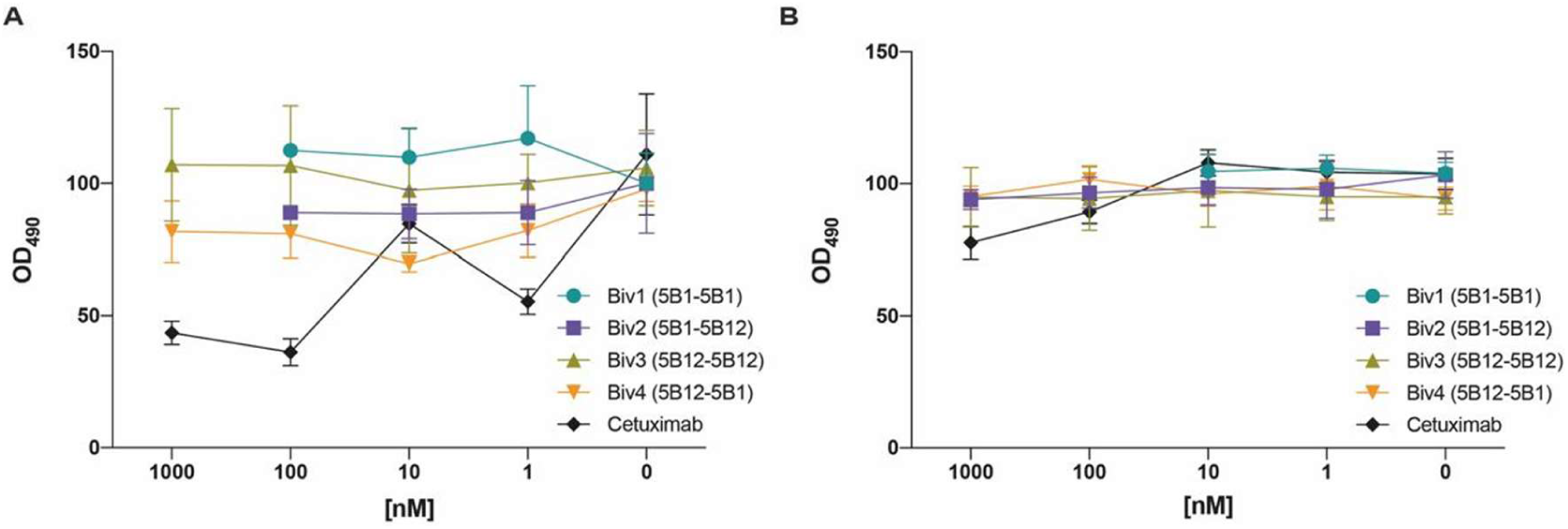
Biparatopic nanobodies inhibit proliferation of glioma cells with the EGFRvIII mutation. Glioma stem cell-enriched cell lines were treated with different equimolar concentrations of the bivalent nanobodies or cetuximab and the inhibition of cell proliferation was measured in triplicates after 6 days by MTS assay. (A) G05 cells expressing the mutated EGFRvIII and (B) G08 cell line expressing non-mutated EGFR.

## 3. Discussion

Nanobodies (Nb) targeting cell surface proteins hold significant promise for therapeutic and diagnostic applications in the cancer field, potentially outperforming mAbs and scFvs. This potential extends beyond their use as stand-alone molecules, as they can also function as modules of Bispecific T-cell Engagers (BiTEs) or for targeting other agents, like CAR-T cells (Yang & Shah, 2020). However, a major limitation to their clinical efficacy, regardless of their form, is the emergence of escape variants that render the therapy ineffective. The epidermal growth factor receptor (EGFR) is a key tumor-associated antigen exploited in cancer therapy (Uribe et al., 2021), but it has been shown to frequently escape antibody-based therapies (Markman et al., 2010). To overcome resistance, strategies such as dual targeting of two tumor-associated antigens (De Munter et al., 2018; Tapia-Galisteo et al., 2022) or the use of antibody mixtures targeting different epitopes (Kearns et al., 2015; Montagut et al., 2018) have been explored. Additionally, biparatopic nanobody fusions have demonstrated the ability to prevent therapeutic failure due to escape variants (Koenig et al., 2021; Walter et al., 2022). In this work, we successfully isolated and characterized novel anti-EGFR Nb. These nanobodies exhibit high affinity for the receptor and can be engineered into bivalent constructs that effectively block EGF-induced EGFR signaling and inhibit EGF-dependent cell proliferation.

To overcome challenges associated with membrane-associated antigen production to target epitopes which are naturally exposed *in vivo*, immunization and selection on cells or alternative tools that present antigens in their native conformation have emerged as valuable approaches (Greenfield et al., 2021). Virus-like particles (VLPs) derived from cells overexpressing the target antigen typically exhibit higher concentration of specific surface antigens compared to whole cells (Tariq et al., 2021), making them more suitable immunogens that significantly enhance the activation of humoral immunity (J. Liu et al., 2022; Tornesello et al., 2022). Therefore, we aimed to generate immune libraries specifically targeting EGFR, expressed on human tumor cells, by immunizing llamas with a combination of EGFR-overexpressing cells and VLPs, allowing us to construct a target-specific library. Initially, when we performed conventional biopanning of this library using well-established methods (Baral et al., 2013; Lipes et al., 2008; Pardon et al., 2014; Peyrassol et al., 2016), it became enriched in defective clones bearing aberrant inserts, a phenomenon previously reported (Carcamo et al., 1998; de Bruin et al., 1999). Insert-less phage clones or clones with premature stop codons are relieved from the metabolic burden of expressing an heterologous protein fused to the gpIII of M13 (de Bruin et al., 1999; Høydahl et al., 2016; Laustsen et al., 2017) which may give them a growth advantage, being preferentially amplified and outcompeting the full-length clones. Limiting the amplification phase after each round of panning should minimize this effect and thus favor the enrichment on target-binding clones. Indeed, we successfully eliminated the selection of defective clones by adjusting this step to 4 h at 28 °C.

While panning with purified proteins is generally a successful and usually straightforward strategy for selecting high affinity Abs against various antigens, it poses several limitations when applied to cell surface antigens (Alfaleh et al., 2017; McGuire et al., 2009; Salema et al., 2016). Immobilization of antigens on solid phase, combined with stringent washing, may alter conformational epitopes that are relevant *in vivo*. Conversely, biopanning on live cells or VLPs has the disadvantage of enriching in phages binding other antigens found on the display surface. Our strategy to isolate EGFR-specific clones involved panning and screening the immune VHH library directly on solid-phase immobilized VLPs or, alternatively, on purified EGFR-Fc bound to magnetic beads. Since the immunization protocol involved the use of both EGFR-overexpressing cells and VLPs, alternating this strategy with the use of purified EGFR ectodomain for biopanning was the most successful method for isolating high-affinity EGFR-binders. Both methods yielded Nb against EGFR, but the efficiency of purified EGFR-Fc selection surpassed that of VLP-selection, successfully yielding high-affinity Nb clones. The impact of different panning strategies on the output of phage-display selection has been described for other libraries (Lou et al., 2001; Lu & Sloan, 1999). The two high-affinity anti-EGFR Nb reported in this study were isolated through panning on protein G-captured EGFR-Fc, but not on passively adsorbed VLP_EGFR_, and their targeted epitopes are present both on native EGFR and recombinant EGFR ectodomain. Positive clones isolated after VLP-based panning showed, in general, lower affinity; this finding is consistent with other studies which showed that selection on high-density displayed antigen, such as on VLPs, can lead to the identification of lower affinity clones due to avidity effects caused by the display of multiple molecules of antigen, resulting in avidity artifacts (Lorimer et al., 1996; Omidfar et al., 2004; Salema et al., 2016).

The two novel high-affinity Nb, 5B1 and 5B12, specifically bind EGFR both in intact cells (as shown by FACS and IF) and as a purified antigen (SPR). Their measured affinities exceed that of the gold standard, 7D12. The equilibrium dissociation constants (K_D_) of 5B1 and 5B12 were of the same order, but with distinct kinetic profiles. 5B12 exhibited a faster association rate, whereas 5B1 displayed a slower dissociation rate. These differences in their kinetic parameters could potentially confer distinct biological capabilities to the nanobodies *in vivo* (Vauquelin, 2016) or even influence the effect in bivalent constructs when they are placed as outer or inner domains (Digiammarino et al., 2011).

Targeting non-overlapping epitopes on EGFR with biparatopic Nb has demonstrated increased anti-tumor activity compared to monospecific nanobodies in preclinical studies (Jin et al., 2013; Roovers et al., 2011). Hence, it is important to map the epitopes targeted by novel Nb. We investigated epitope overlap by competitive ELISA. Our results show that 5B1 competes with cetuximab for binding to EGFR, whereas 5B12 could be targeting partially overlapping or separate but allosteric epitopes.

It has been previously shown that inhibition of EGF-driven proliferation by anti-EGFR Abs depends on bivalence, as the mechanism involves dimerization and down-regulation of the receptor (Fan et al., 1994; Harwood et al., 2017). In fact, our results show that monovalent Nb poorly inhibited A431 proliferation, whereas bivalent Nb were almost as efficient as cetuximab. Nb were also assayed for their capacity to inhibit EGF-induced EGFR phosphorylation; only bivalent constructs inhibited the EGF-induced phosphorylation of tyrosine 1068 (which has been shown to be an early event in signaling towards Ras (Buday & Downward, 1993)).

Moreover, the bivalent, biparatopic constructs were able to inhibit the in vitro proliferation of patient-derived glioma cells expressing the mutated truncated EGFRvIII form. Therefore, we could consider that the significant anti-proliferative properties may be related to the functional bivalent conditions of these molecules. Surprisingly, the biparatopic nanobodies that were able to inhibit cell proliferation and blocked EGFR phosphorylation, showed no increased affinity compared to the monovalent nanobodies. Previous studies have shown that the therapeutic efficacy of biparatopic Abs often arises from combining paratopes with distinct yet complementary mechanisms of action. For example, biparatopic molecules may achieve enhanced potency through induced fit, rapid rebinding, additional steric hindrance due to their increased size, or conformational changes that amplify receptor inhibition (Beirnaert et al., 2017; Hultberg et al., 2011; Ma et al., 2021; Roovers et al., 2011). Thus, the ability of bivalent constructs to inhibit EGF-induced phosphorylation and cell proliferation, despite not exhibiting higher affinity than their monovalent counterparts, may be attributed to alternative mechanisms of action.

In conclusion, in this study we reported the isolation of two novel high-affinity anti-EGFR nanobodies, which target different epitopes and are amenable of construction of biparatopic nanobodies with improved properties. These mono and bivalent constructs could become useful tools for the diagnosis and treatment of EGFR-associated tumors.

## 4. Materials and Methods

### 4.1. Cell lines

HEK293A cells were cultured in DMEM high glucose medium supplemented with 10% fetal bovine serum (FBS) and 1% penicillin-streptomycin. The his-tagged full-length human EGFR sequence (Cat#HG10001-C-H, Sino Biological, Wayne, PA, USA) was subcloned downstream of a CMV promoter in a pCMV expression plasmid with an hygromycin resistance cassette (Sino Biological, Wayne, PA, USA). HEK293A cells were transfected and individual hygromycin resistant clones were established. An EGFR-expressing HEK293A cell clone (HEK293A_hEGFR_) was selected for llama immunization following western blot analysis after sample solubilization in 2X Laemmli sample buffer (65.8 mM Tris-HCl, pH 6.8, 2.1% SDS, 26.3% (w/v) glycerol, 0.01% bromophenol blue) using QIAexpress anti-His antibodies (Qiagen, Hilden, Germany) and a rabbit monoclonal antibody against EGFR (Cat#10001-R021, Sino Biological, Wayne, PA, USA). EGFR expression levels on the cellular surface were quantified using the Quantum^TM^ Simply Cellular kit (Bangs Laboratories Inc., Fishers, IN, USA, Cat#814) using anti EGFR-PE as specified by the manufacturer.

A431 or MCF7 cells were cultured in DMEM high glucose or DMEM-F12 respectively supplemented with 10% FBS and 1% penicillin-streptomycin.

### 4.2. Virus-Like Particle production

The EGFR virus-like particles (VLPs) were pseudotyped murine leukemia virus (MLV) carrying the full length, membrane integral EGF receptor (Balliet & Bates, 1998). VLPs were produced as previously describe by co-transfection of HEK293 cells with pCGP encoding the structural (gag) and (pol) genes of MLV (named p177) and either p2231, encoding EGFR to generate EGFR-VLPs, or pBSII null plasmid to generate control, ‘naked’ or null-VLPs as previously described (Hoffman et al., 2000; Rucker, 2003). Modifications of the previous method included using polyethylenimine (PEI) as the transfection agent in place of calcium-phosphate and testing co-plasmids mixtures to attain the highest EGFR levels on the VLPs, typically at ratios of 3 to 5 receptor per pCGP. VLPs were harvested twice, by collecting the supernatant at 48 h after transfection, by replacing it with fresh DMEM-5% FBS, then incubating for an additional 24-48 h before a second collection. Both supernatants were filtered through a 0.45 µm filter and concentrated by pelleting through 20% sucrose/HEPES at 140,000 g for 1 h at 4°C. The pellets were re-suspended in 10 mM HEPES pH 7.5 and stored at 4°C as described (Balliet & Bates, 1998; Hoffman et al., 2000; Rucker, 2003). The protein content of VLPs was quantified using the bicinchoninic acid (BCA) titration assay (Cat#23227, Waltham, MA) according to manufacturer’s instructions. The success of the co-transfection of VLPs with EGFR was assessed by dot-blot and the EGFR-VLPs quantification by VLP-ELISA to attain signal saturation. Both methods used anti-CD81 (Cat#MA5-13548, Carlsbad, CA), a HEK293 cell surface marker and anti-EGFR (Cat#10001-R021, Sino Biological, Wayne, PA) antibodies at the recommended dilution as reporters. The plasmids p177, p2231, and pBSII were a gift from Drs. Willis and Doranz (Integral Molecular, Philadelphia, PA).

### 4.3. Llama immunization

To induce a humoral immune response directed to EGFR, two llamas were injected subcutaneously with approximately 2 x 10^7^ HEK293A_hEGFR_ cells and boosted twice every two weeks with 10^7^ HEK293A_hEGFR_ cells. A final boost with 70 U VLP_EGFR_ was administered six weeks after the prime. Pre-immune and immune sera were collected at days 0, 35 and 49 and the specific antibodies were monitored by ELISA. Four and seven days after the last antigen injection, two 100 ml blood samples were collected and peripheral blood lymphocytes (PBLs) were purified by density gradient centrifugation on Ficoll-Paque gradients (Amersham Biosciences, Little Chalfont, UK).

All animal experiments were conducted with the approval of Fundación Instituto Leloir Institutional Animal Care and Use Committee.

### 4.4. Phage library construction and selection

Phage-displayed Nb libraries were constructed from the peripheral blood B lymphocytes of the immunized llamas as described previously (Fig. 6) (Alzogaray et al., 2019). Briefly, mononuclear cells were isolated from 100 ml of blood by Ficoll-Paque (GE Healthcare, Chicago, USA) gradient centrifugation. Total RNA was purified from these cells using TRIzol (Invitrogen, Carlsbad, CA, USA) and cDNA was reverse transcribed. VHH genes were amplified using semi-nested PCR. The amplicons were purified from agarose gels, digested with SfiI and NotI (New England Biolabs, Ipswich, MA, USA) and cloned into the pHEN2 phagemid vector downstream the PelB-leader peptide and upstream the chimeric His6x-Myc epitope tag. Transformation into *Escherichia coli* XL1-Blue yielded libraries with a size of approximately 10^7^ clones independent transformants and an insert rate of ∼89% as assessed by colony PCR of 19 clones. Ten clones were used for plasmid DNA preparation (QIAprep Spin Miniprep Kit, Qiagen, Hilden, Germany) and sequenced for library diversity analysis (Macrogen, Seoul, Korea).

### 4.5. Screening of EGFR-specific Nb by phage display technology and phage ELISA identification

The nanobodies were selected by phage display. Phage particles were rescued from the library using M13K07 helper phage (Life Technologies, Carlsbad, USA) and panned by two alternative methods. For panning on plate-adsorbed VLPs, wells of NUNC® MaxiSorp™ microtiter plates (Thermo Fisher Scientific, Waltham, MA, USA, Cat#476635) were coated overnight at 4 °C with 100µl/well of VLP_EGFR_ or VLP_null_ (100 ng/well in carbonate buffer pH 9.6). The plates were washed with 0.05% Tween 20-PBS, and then were blocked with 200 µl of 5% skim milk in PBS pH 7.4 for 1 h at 25 °C. After another wash with 0.05% Tween 20-PBS phages from each library were added to the plates and incubated for 1 h at room temperature. After incubation, the plates were washed with 0.05% Tween 20-PBS and bound phage particles were eluted with 100 mM triethylamine (TEA, pH 10.0) and immediately neutralized with 1M Tris (pH 7.4). The eluted phages were used to infect exponentially growing *E. coli* TG1. For panning on the soluble purified EGFR ectodomain (ECD) fused to the Fc portion of IgG, EGFR-Fc (Sino Biological, Wayne, PA, USA, Cat#10001-H02H), was captured on Protein G magnetic beads (New England BioLabs, Ipswich, MA, USA, Cat#S1430S). Phages were incubated with 25 µL EGFR-magnetic beads complex at room temperature in a rotator for 2 h to mix them mildly. The magnetic beads were separated by a magnetic rack to wash off the phages which did not bind to the magnetic beads. The magnetic beads were resuspended and eluted with 100 mM TEA (pH 10.0) solution and immediately neutralized with 1M Tris (pH 7.4). The biopanning protocol was followed as already described for the solid phase strategy.

The eluted phages were used to infect *E. coli* TG1 in 30 ml of 2 × YT medium, and the cultures were allowed to grow during the exponential phase until they reached OD_600_ = 0.5, when the M13K07 helper phages were added to rescue the phages. After infection, samples were incubated ON at 37 °C, 200 rpm. The above steps describe one round of selection, and these rescued phages were used in the next round of selection. After three rounds, the eluted phages were used to infect *E. coli* TG1 cells. Ninety individual colonies were randomly selected and picked into 96-well plates containing 200 µl of 2 × YT medium overnight.

### 4.6. Monoclonal phage ELISA

*E. coli* TG1 bacteria were infected with the eluted phage for 20 min at room temperature. Subsequently, the infected cells were spread evenly on LB + ampicillin plates. Following an ON incubation period at 37 °C, colonies of *E. coli* TG1 were chosen at random and inoculated into 200 µl of LB + ampicillin in sterile 96-deep-well plates (Corning, NY, USA). Individual *E. coli* phage clones were superinfected with M13KO7 at 28 °C ON. To perform the phage ELISA, 100 microliters of culture supernatant containing phage particles from the 96-well plates were applied to microtiter plates according to the specified method (Pardon et al., 2014). In order to establish the negative controls, BSA was utilized as an antigen negative control, while M13K07 helper phage served as the phage negative control. Color development was performed by the addition of 50 µL of TMB Single Solution (Thermo Fisher Scientific, Carlsbad, CA, USA). After 8 min, the enzyme reaction was stopped with 50 µL of 1 M sulfuric acid per well, and the absorbance was measured in a Bio-Rad Model 550 microplate reader (Bio-Rad Laboratories, Hercules, CA, USA) at 450 nm.

### 4.7. Competitive ELISA

Wells of NUNC® MaxiSorp™ microtiter plates were coated overnight at 4 °C with 100 µL of VLP_EGFR_. The wells were blocked with 200 µl of PBS containing 5% skim milk for 1 h at RT. Serially diluted cetuximab (0.007µM to 50µM, Erbitux® CETUXIMAB, MERCK) and a fixed amount of VHH phage was added to wells for 1 h at room temperature. The wells were washed 5 times with PBS containing 0.1% Tween-20, incubated for 1 h with anti-M13 conjugated with Horseradish Peroxidase (HRP), washed 5 times again and then developed with TMB substrate. The absorbance was measured using an ELISA reader at 450 nm. The competitive activity of Nb was expressed as a percentage of displacement of cetuximab ((1-((OD_450_ of sample-OD_450_ of blank) / (OD_450_ of Nb without cetuximab-OD_450_ of blank)))/x 100 %).

### 4.8. Expression and purification of Nb

VHH DNA sequences coding for 5B1 and 5B12 (Fig. S5) were cloned into the pHEN6 expression vector and the recombinant plasmids were transformed into *E. coli* WK6. The transformed cells were cultured at 37 °C and 180 rpm in LB medium containing 100 µg/ml ampicillin until they reached the logarithmic growth phase (OD_600_ = 0.4–0.8). IPTG was added to a final concentration of 1 mM, and the cells were then cultured at 28 °C and 180 rpm ON. The cultures were centrifuged at 10,000 g for 10 min at 4 °C, cells were frozen at −80 °C and thawed at RT, and resuspended in TES buffer (100 mM Tris–HCl pH 8.0, 1 mM EDTA, and 20% sucrose). The supernatant corresponding to the periplasm containing VHHs was collected by centrifugation at 10,000 g for 30 min at 4 °C and dialyzed against equilibration buffer (50mM NaH_2_PO_4_, 300mM NaCl, 10 mM imidazole, H_2_O_dd_, pH 8) ON at 4 °C using a 3,500 Da cut-off dialysis cellulose membrane (Spectrum™ Spectra/Por™ Dialysis Membrane Tubing, MWCO 3.5KD, ThermoFisher Scientific, Waltham, MA, USA). Monomeric VHHs tagged C-terminally with a 6×His tag were purified from the periplasmic extract of *E. coli* WK6 cells by immobilized metal affinity chromatography (IMAC) using a His Trap^TM^ HP column (GE Healthcare, Chicago, USA).

7D12 is a public domain antibody and its sequence was obtained from Gainkam et al. (Gainkam et al., 2008) and synthesized by Genscript (Pistacaway, NJ,USA). 7D12 was cloned into pHEN6 and expressed and purified as the rest of the Nbs.

### 4.9. Immunofluorescence

HEK293A_hEGFR_ or HEK293A cell lines were grown on a glass coverslip, washed with PBS and fixed with 4% paraformaldehyde for 30 min. The samples were blocked with 10% FBS/PBS for 1 h. Samples were incubated with the VHHs (500nM) ON in PBS/2.5% FBS, followed by anti-HIS-TAG antibody (Abgent, San Diego, CA, USA) for 1 h, and anti-mouse Cy5-conjugated secondary antibody (Jackson ImmunoResearch, PA, USA) for 1 h. Cells were washed four times with PBS and mounted on glass slides. All samples were labeled with DAPI (Sigma-Aldrich, San Luis, USA) and phalloidin (Sigma-Aldrich, San Luis, USA).

### 4.10. Flow cytometry analysis (FACS)

A431 or MCF7 cells were harvested with Accutase and 5 × 10^5^ cells were incubated with different concentrations of nanobodies in 500 µl of 3% BSA PBS for 2 h at 4 °C in head over tail rotation. Anti-BLS was used as a negative control. The cells were washed three times with 500 µl of 3% BSA PBS at 4 °C to remove unbound antibody. The cells were resuspended and incubated with anti-His-Tag antibody (1:2000). After incubation, the cells were washed three times with 500 µL of 3% BSA PBS, and resuspended in 500 µL of 3% BSA PBS. The cells were incubated with a Cy5-conjugated mouse anti-His antibody for 30 min at 4 °C in 100 µl of 2% (w/v) BSA in PBS, and were washed three times with 500 µl of 3% BSA PBS to remove unbound antibody. The samples were acquired using a FACS Aria flow cytometer (Beckman Coulter, Brea, CA, USA) and the data analyzed using the FlowJo™ v10 software (BD Biosciences, San Jose, CA).

### 4.11. Studies to assess cell viability

A431 cells plated at 5,000 cells/well in 96-well plates were incubated with DMEM containing 10% (v/v) FBS for 6 h, followed by incubation with FBS-free DMEM for 24 h. The medium was replaced with the respective Nb or Cetuximab at the appropriate concentration and 170 pM EGF. After incubation for 48 h at 37 °C 5% CO_2_, cells were treated with MTS (CellTiter 96 Aqueous One Solution Cell Proliferation assay, Promega, Madison, WI, USA) at 5 mg/ml for 2 h at 37 °C 5% CO_2_. Optical density of the wells was read at 490 nm on a plate reader (iMark™ Microplate Absorbance Reader, Bio-Rad, Hercules, CA, USA). The viability of the cells was calculated as a percentage relative to the control wells treated with media alone (100% viability). All the experiments were performed in triplicate.

### 4.12. Binding Analysis by Surface Plasmon Resonance (SPR)

The interaction between the different Nb and EGFR was studied using surface plasmon resonance (SPR) with a BIAcore T100 instrument (Cytiva, Marlborough, MA, USA). EGFR-Fc (the ligand) was diluted in sodium acetate pH 4.5 to a final concentration of 15 µg/ml and immobilized (500 RU) on a CM5 chip surface (Cytiva, Marlborough, MA, USA) through amine coupling, following the manufacturer’s instructions. Soluble VHHs were diluted in PBS, pH 7.4 supplemented with 0.005% Tween as the running buffer and injected over the chip surfaces at a flow rate of 50 µl/min for at least 90 sec at 25 °C. The data were analyzed using the BIA evaluation software (Cytiva, Marlborough, MA, USA) considering analyte concentration in the range 100-1.56 nM. The dissociation was monitored for 120 s and glycine pH 3.0 was injected on the surface for 30 s for its regeneration. PBS was not able to wash off the VHHs after interaction with the ligand. The data represent the average of at least two independent assays for VHHs 5B1, 5B12, and 7D12 under each condition. The dissociation constant (K_D_) of the system was determined from the kinetic parameters, the association and dissociation rate (*k_on_, k_off_*) by a kinetic analysis using a 1:1 binding model when it was possible. The bivalent nanobodies did not allow this analysis and the K_Ds_ were determined under state steady affinity conditions (SSA method). The standard deviation (SE) of each result was obtained using the BIA evaluation software.

### 4.13. Statistical analysis

All statistical analysis, calculations and tests were performed using GraphPad Prism 8 (v 8.2.0, GraphPad Software, San Diego, CA) and FlowJo™ Software (v 10, BD Biosciences, San Jose, CA).

## 5. Conclusions

Using two alternative approaches aiming to isolate sdAb recognizing EGFR in its native conformation, we were able to isolate and characterize two novel nanobodies. We compared the characteristics of these nanobodies with the publicly available Nb 7D12 and the commercially available anti-EGFR mAb cetuximab. Remarkably, both 5B1 and 5B12 nanobodies exhibited higher affinity for EGFR than 7D12. Furthermore, bivalent constructs of these nanobodies demonstrated to be effective in the inhibition of malignant cell growth. Given these compelling findings, it is evident that the novel nanobodies, whether in their monovalent or bivalent configurations, hold significant promise as prospective candidates for innovative applications in targeting EGFR.

## 6. Patents

SEV and OLP and co-inventors of a related provisional patent application assigned to Fundación Instituto Leloir and CONICET.

## Author Contributions

Conceptualization, O.L.P and S.E.V; methodology, M.H., M.M.F., V.Z., S.E.V.; formal analysis, M.M.F., E.L.M. and S.E.V.; investigation, M.H., M.M.F., D.C.A.-C., S.W., G.C. and V.Z.; writing—original draft preparation, M.M.F., O.L.P. and S.E.V.; writing—review and editing, M.H., M.M.F., E.L.M., O.L.P. and S.E.V.; visualization, M.H., M.M.F., D.C.A.-C. and S.E.V.; supervision, O.L.P and S.E.V.; project administration, S.E.V.; funding acquisition, O.L.P. All authors have read and agreed to the published version of the manuscript.

## Funding

This research has been funded in part by the Argentinean National Agency of Research, Development and Innovation through an EMPRETECNO grant of Fonarsec; we acknowledge the continuous support of the organization Amigos de Instituto Leloir para la Investigación en Cáncer (AFULIC), Rio Cuarto, Argentina.

## Institutional Review Board Statement

Not applicable.

## Informed Consent Statement

Not applicable.

## Data Availability Statement

Data will be provided by request.

## Acknowledgments

We deeply thank Dr. Vanina Alzogaray for anti-BLS Nb and helpful discussions, Soledad Malori for her technical assistance with the chromatography experiments and Dr. Carla Pascuale for her technical assistance with the flow cytometry experiments.

## Conflicts of Interest

The authors declare no other conflict of interest. The funders had no role in the design of the study, the collection, analyses, or interpretation of data, the writing of the manuscript or in the decision to publish the results.

## Notes

### Competing Interest Statement

The authors have declared no competing interest.

