## Supplemental Figures and Tables for "Development of novel high-affinity nanobodies against EGFR for cancer therapy"

### Supplementary Material

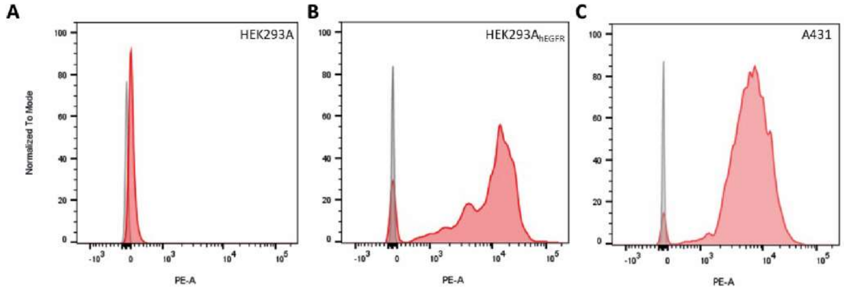


D

| Sample | ABC | MFI |
|---|---|---|
| HEK293A | 128 | 114 |
| HEK293A$_{hEGFR}$ | 81338 | 6874 |
| A431 | 70597 | 6283 |

**Fig. S1: Determination of EGFR Density.** (A) HEK293A (B) HEK293$_{hEGFR}$ and (C) A431 cells were labeled with a monovalent anti-EGFR antibody using the QuantumTM Simply Cellular kit, signal was detected using BD FACSAria™ Cell Sorter and data were analyzed using the FlowJo™ v10 software (BD Biosciences). Labeled cells are shown in red and non-labeled control cells in grey. (D) ABC (Antibody Binding Capacity) value is equivalent to the number of cell surface receptors per cell.

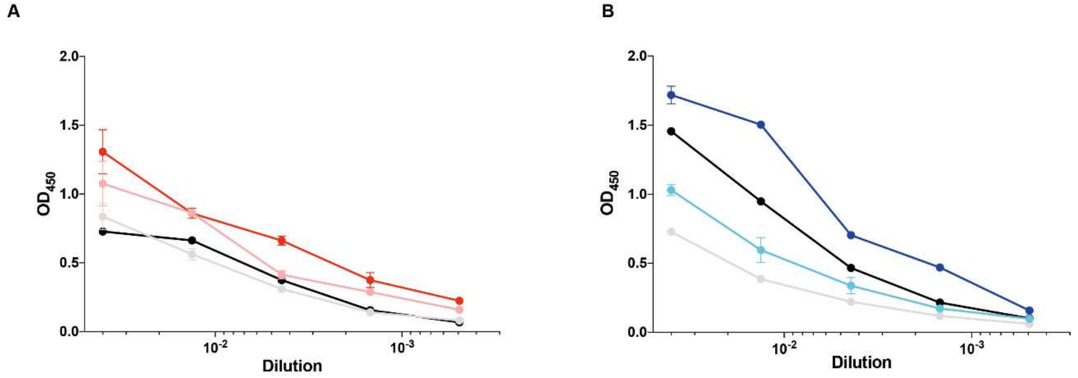


**Fig. S2: Immunization of *Llama glama* with EGFR-overexpressing cells and VLPs induces an anti-EGFR humoral immune response.** The reactivity of pre-immune (day 0: grey/pink/cyan) and immune sera (day 35: black/red/blue) of the llama was assessed by ELISA against purified antigens (A) or VLPs (B). For protein-coated ELISA the negative control was BSA (grey/black) and the specific antigen was EGFR-Fc (pink/red). For VLP-coated ELISA the negative control was $VLP_{null}$ (grey/black) and the specific antigen was $VLP_{EGFR}$ (cyan/blue).

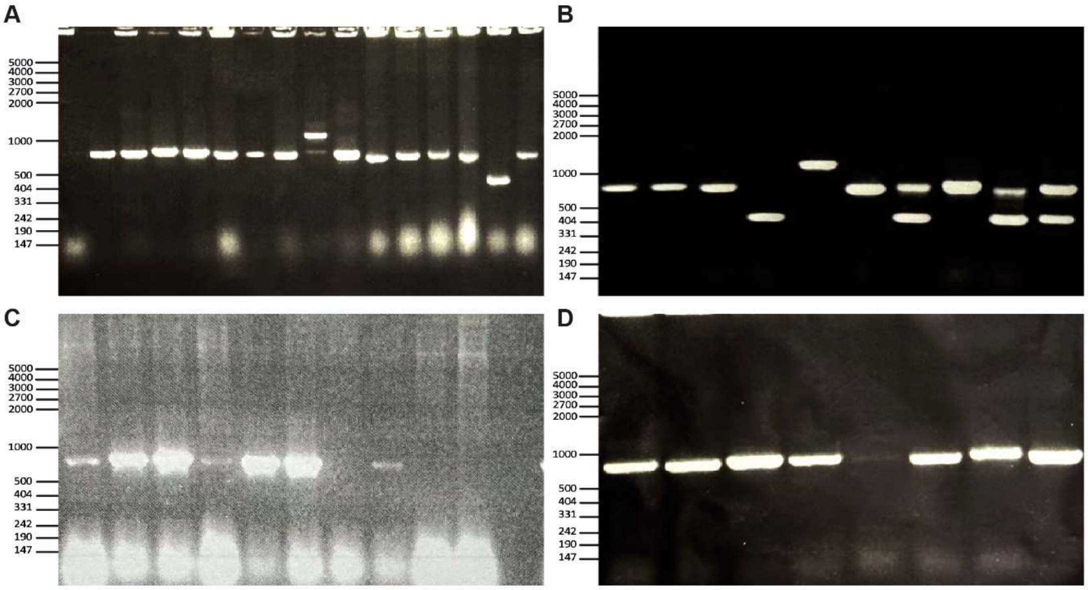


**Fig S3. Colony PCR of clones after each biopanning round.** Ten to 15 clones were picked at random to determine the percentage of full size VHH sequences in each round of biopanning of phage-displayed VHHs (A) Colony-PCR of the library before panning, where the full-length inserts represent 86.7% of the total clones. (B) After the first round of selection and amplification full-length inserts represent between 50-70% of the total. After the second (C) and third (D) round, the percentage of full-length inserts was found to be between 90-100%.

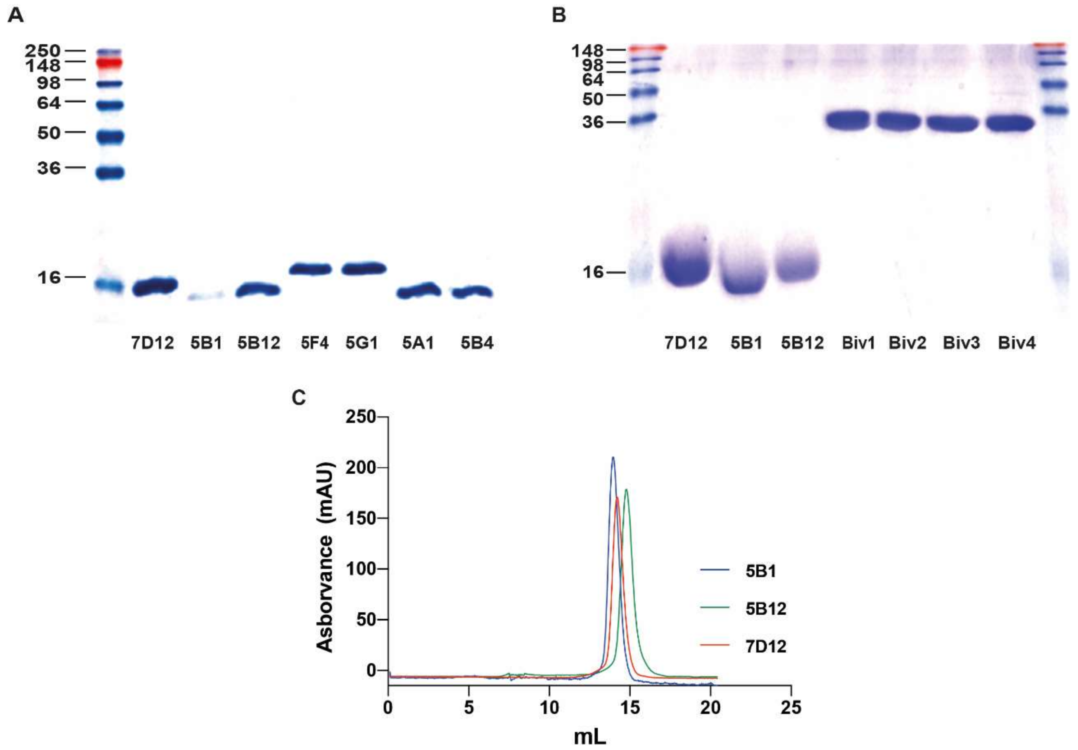


**Fig S4. Expression and purification of Nb.** (A) SDS-PAGE of His-tagged purified Nb. (B) SDS-PAGE showing the correct expression of the corresponding bivalent constructs (~30 kDa). (C) Size-exclusion chromatography profile of the IMAC-purified 7D12, 5B1 and 5B12; nanobodies were concentrated on Vivaspin concentrators with 5 kD molecular weight cut-off (Sartorius Stedim Biotech GmbH, Goettingen, Germany) and loaded onto a superdex S75 (SuperdexTM 75 10/300 GL) column in PBS buffer.

**>5B1**
QVQLQESGGGLVQAGDSLRLSCAASGRTFSTYTMGWFRQAPGKEREFVAGIGYSDDNTYYTDSVK
GRFTVSRGSAKNTVYLQMNSLKPEDTAVYYCAARVGDVVFTIADNYAYWGQGTQVTVSS

**>5B12**
QVQLQESGGGLVQTGDSLRLSCAASGGTLTSYNMGWFRQAPGKDREFVAGISWSGGTGTYYADSV
KGRFTISTDNAKNTVYLQMDSLKPDDTAVYYCAVRRRAYSVYTRPGLYDYWGQGTQVTVSS

**Fig S5. Nanobody sequences.**

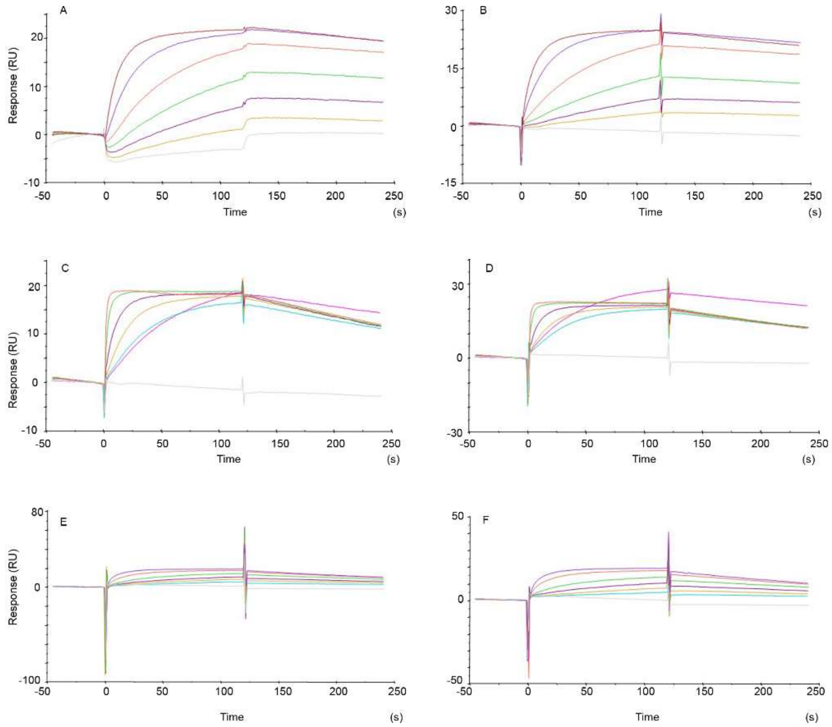


**FigS6. Surface plasmon resonance analysis of the interaction between VHH and EGFR at a flow rate of 50 µl/min.** SPR sensorgrams showing the interaction of immobilized EGFR and soluble VHHs. (A, B) 5B1 (100-3.125 nM); (C,D) 5B12 (100-3.125 nM); (E, F) 7D12 (100-3.125 nM).

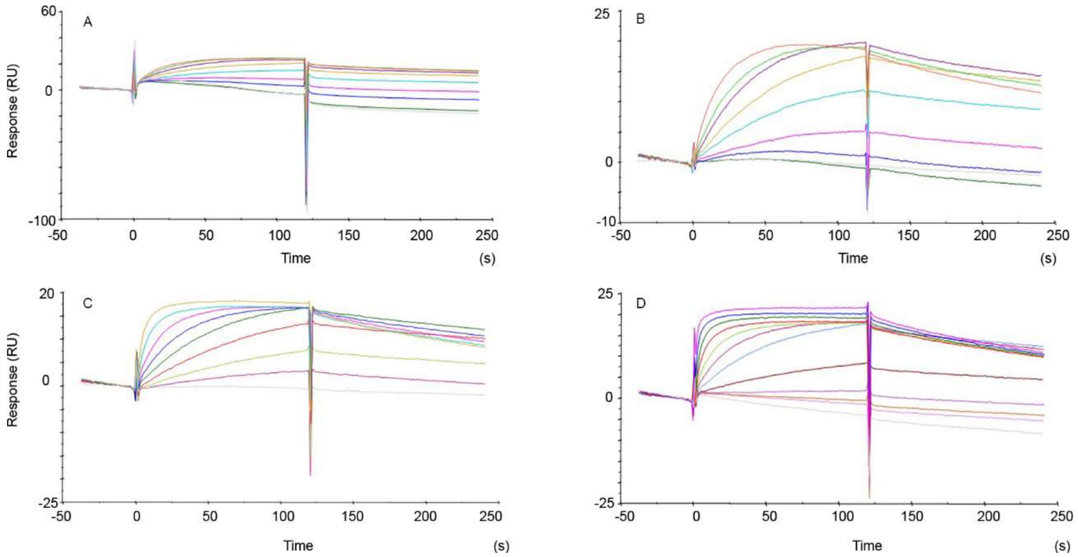


**FigS7. Surface plasmon resonance analysis of the interaction between VHH and EGFR.** SPR sensorgrams showing the interaction of immobilized EGFR and soluble VHHs at a flow rate of 50 μl/min. (A) 5B1-5B1 (100-0.8 nM); (B) 5B1-5B12 (200-0.8 nM); (C) 5B12-5B12 (100-0.8 nM); (D) 5B12-5B1 (800-0.8 nM).

**Table S1: Kinetic parameters of the interaction between EGFR and VHHs determined at a flow rate of 50 μl/min.**

| Analyte[1] | $K_{on}$ (M.s)$^{-1}$ | $K_{off}$ (s$^{-1}$) | $K_D$ (M) |
|---|---|---|---|
| 5B1 | (7.4 ± 0.1) $10^5$ | (1.14 ± 0.02) $10^{-3}$ | 1.6 x $10^{-9}$ |
| | (1.72 ± 0.01) $10^6$ | (4.5 ± 0.4) $10^{-4}$ | 5.9 x $10^{-10}$ |
| 5B12 | (6.5 ± 0.3) $10^6$ | (7.7 ± 0.4) $10^{-3}$ | 5.6 x $10^{-10}$ |
| | (3.4 ± 0.3) $10^7$ | (3.65 ± 0.09) $10^{-3}$ | 1.4 x $10^{-10}$ |
| 7D12 | (9.4 ± 0.4) $10^5$ | (2.9 ± 0.3) $10^{-3}$ | 3.0 x $10^{-9}$ |
| | (8.0 ± 0.2) $10^5$ | (3.4 ± 0.2) $10^{-3}$ | 3.9 x $10^{-9}$ |

[1] Monovalent Nb were assayed in two independent assays.
